# State-dependent mapping of GlyR-cholesterol interactions by coupling crosslinking with mass spectrometry

**DOI:** 10.1101/2020.07.02.185280

**Authors:** Nicholas A. Ferraro, Michael Cascio

## Abstract

Pentameric ligand-gated ion channel (pLGIC) allostery is dependent on dynamic associations with its diverse environment. The cellular membrane’s lipid composition influences channel function with cholesterol being a key regulator of channel activity. Human **α**1 glycine receptor (GlyR) was purified from baculovirus infected insect cells and reconstituted in unilamellar vesicles at physiological cholesterol:lipid ratios with aliquots of azi-cholesterol, a photoactivatable non-specific crosslinker. The receptor in vesicles was then enriched in either a resting, open, or desensitized state prior to photocrosslinking. Following photoactivation, crosslinked cholesterol-GlyR was trypsinized and sites of direct covalent attachment to peptides were identified by targeted MS/MS. Dozens of state-dependent crosslinks were identified and differential patterns of cholesterol-GlyR crosslinks were observed in the extracellular region nearing the lipid bilayer, in the M4 transmembrane helix, and in the large intracellular M3-M4 loop. Unique crosslinks in comparative studies identify changes in lipid accessibility or modulation of hydrophobic cavities in GlyR as a function of receptor allostery. Most notably, the outward twisting of M4 and differential crosslinking within the M3-M4 loop provide new insight into allosteric repositioning of GlyR. More generally, this study provides an accurate and sensitive approach to mapping the protein-lipid interactions to discern state-dependent structural movements of membrane proteins embedded in lipid-bilayers.

**Significance:** Ion channels are highly allosteric molecular machines whose structure and function are sensitive to lipids and ligands. While the structures of many pLGICs are known, these are often truncated forms of the receptor in a membrane-mimetic environment locked in ligand-bound conformational states that may not accurately reflect the conformation and dynamics of the receptor in a native lipid environment. Crosslinking coupled with mass spectrometry (CX-MS) has the capability of interrogating the structure of full-length receptors in a lipid environment. In this study, CX-MS was used to identify state-dependent cholesterol-GlyR interactions to identify differential cholesterol accessibility as a function of channel dynamics upon gating and desensitization.

## Introduction

Protein function is modulated by dynamic interactions with other biomolecules such as metabolites, proteins and lipids, and the cellular membrane’s lipid composition has been increasingly recognized as a major contributor in membrane protein function^1,2^. The current model of the lipid membrane suggests a non-homogenous distribution of lipids and proteins that form microdomains, with the lipid composition modulating cellular processes at the protein-lipid interface either directly (i.e., binding) or indirectly (altering the physiochemical properties of the bilayer)^3–5^. Examples of these effects are the influence of lipid composition on the structure/stability of the transmembrane domain of amyloid precursor protein^6^, the modulation of β2-adrenergic receptor dimer interface stabilization through cholesterol occupancy^7^, the abrogation of dimer formation of the leucine transporter from cardiolipin delipidation^8^, and the effects of cholesterol on ion channels^9–11^. Lipids modulate agonist binding of glutamate-gated chloride channel (GluCl) through occupancy of membrane-spanning intersubunit crevices, promoting an expanded, open-like conformation that potentiates the receptor^12^. The potency of γ-aminobutyric acid receptor (GABAR) is diminished by cholesterol depletion and restored through cholesterol enrichment of neurons^13^. Lipids are frequently observed co-crystallized with membrane proteins, underlining the specificity of certain protein-lipid interactions and implying functional roles in phsyiology^14^. For example, cholesterol co-crystallizes with G-protein coupled receptors^15^, Na^+^,K^+^ ATPase^16^, and the dopamine transporter^17^, and phospholipid co-crystallizes with GluCl^12^ and *Gloeobacter* ligand-gated ion channel (GLIC)^18^.

Many pharmaceuticals (anesthetics, barbiturates, benzodiazepines, cannabinoids, and alcohol) and bioactive lipids (progesterone, sphingomyelin, and ceramide) specifically target the pLGIC superfamily of ion channels^19–22^. Anesthetics alter the permeability of both anion-selective and cation-selective channels. However the molecular mechanism of channel modulation remains poorly understood^23^. Bioactive lipids play a diverse role, including the regulation of cell-surface nicotinic acetylcholine receptor (nAChR) levels^21^, the inhibition of GABAR^20^, and the modulation of nAChR desensitization^24^. Intriguingly, both anesthetics and bioactive lipids partition at the lipid-protein interface in cellular membranes^25–27^, with the former being observed in crystal structures of pLGIC structural homologues^28,29^. Cannabinoids such as Δ^9^-tetrahydrocannabinol potentiate GlyR activity via interactions at the protein-lipid interface^30^. Screening of GlyR for cannabinoid-like potentiating agents is an effective tool to discover novel therapeutics^22^ highlighting the importance in developing methodologies to sensitively probe protein-lipid interactions. Direct effect of anesthetic agents and alcohol of pLGICs provide valuable models for general allosteric modulation and to further develop anesthetic agents^31^.

Cholesterol and saturated lipids enrich lipid rafts forming highly ordered microdomain complexes distinct from the surrounding disordered lipid environment, with this highly ordered domain providing a mechanism for protein interactions and the regulation of cellular processes^32^. Cholesterol in molar excess of the capacity of these complexes has high fugacity, and is regarded as “active cholesterol”^33^, as seen by an abrupt increase in sterol availability to cholesterol oxidase^34^, perfringolysin^35^, and methyl-β-cyclodextrin^36^ emerging at concentrations above 25-35 mol percent^37,38^. Excess cholesterol exhibits high chemical activity in a chemical phase distinct from that observed under negligible chemical activity^39^ that may potentially drive regulatory processes within or on the plasma membrane surface^40^, either indirectly by modulating plasma membrane physical properties or directly as a protein regulator^41^. Cholesterol and anionic phospholipids modulate nicotinic acetylcholine receptor (nAChR) allosteric transitions whereby cholesterol enrichment of cholesterol-depleted membranes up to a given threshold (∼35 mol%) enhanced receptor-mediated ion flux from inactive channels to being able to undergo agonist-induced state transitions^42–46^. This profound regulatory effect of cholesterol also causes nAChR to adopt distinct conformations as a function of cholesterol concentration, where in the absence of cholesterol or anionic phospholipids adopting a conformation that has properties distinct of the resting or desensitized state in which the allosteric coupling between neurotransmitter binding sites and the transmembrane pore is lost^47,48^. Similarly, crosslinking-mass spectrometry (CX-MS) studies identified differential GlyR-cholesterol crosslinking patterns at low and high cholesterol concentrations, suggesting that GlyR adopted distinct conformations as a function of cholesterol concentration^49^.

The temporal quality of neuronal receptor allostery typically poses a challenge for experimental characterization as some allosteric states exist transiently, however the approach of using photocrosslinking provides an opportunity to probe the dynamics of receptor gating and desensitization^50,51^. Photoaffinity labeling studies identified the propofol-binding site of GLIC^52^ and GABAR^53^ as well as cholesterol interactions with the peripheral-type benzodiazepine receptor^54^, nAChR^55,56^, and GlyR^49^. The incorporation of sensitive mass-spectrometry (MS)-based approaches to sensitively identify sites of photocrosslinking has the potential to identify allosteric dynamics of protein structure in membranes and to examine the role of lipids in receptor allostery^49,51^. Current tandem MS instruments have the mass accuracy and sensitivity to unambiguously identify and refine crosslinked species to amino acid residue(s) within a given peptide^49,57,58^. CX-MS studies provide a valuable adjunct to crystallography and cryo-EM studies particularly in less resolved regions of images by providing amino acid proximity and distance constraint information useful in homology or *de novo* modeling^59–62^. Data provided by state-dependent CX-MS can supplement assays elucidating channel mechanisms of activation/desensitization^63^ and allosteric coupling of domains^64^ by providing dynamic localized information regarding structural changes often in unresolved regions of proteins.

Given the essential, yet poorly characterized structural effects of cholesterol modulation of pLGICs, we examined cholesterol accessibility of GlyR at physiological levels, >40 mol% (above its activity threshold), in a state-dependent manner. This study focuses on comparative cholesterol-GlyR interactions, i.e. comparing differential azi-cholesterol crosslinking under elevated “active” cholesterol conditions of the resting, open (F207A/A288G, 30 nM ivermectin^65^), and desensitized state (10 mM glycine) using photocrosslinking coupled with multidimensional MS. The F207A/A288G double mutant GlyR produces an ivermectin-sensitive channel unable to desensitize^65^. These studies expand upon a previous study^49^ that contrasted azi-cholesterol crosslinking to the resting state of α1 GlyR under low and high cholesterol levels. GlyR in the presence of saturating concentrations of glycine activate and stabilize channels predominantly in a higher-affinity ligand-bound conformation, with potential sampling of other states. F207A/A288G GlyR in the presence of nanomolar concentrations of ivermectin has been shown to stabilize channels in a conducting conformation that does not desensitize^65^. Azi-cholesterol, a photoactivatable cholesterol analog, was crosslinked to human homomeric α1 GlyR (wild-type and F207A/A288G) at conditions that probe interactions when cholesterol is chemically active (>40 mol percent natural cholesterol in membrane) in a state-dependent manner^39,66^. This study shows that photocrosslinking coupled with tandem MS can map a lipid-protein interface in several allosteric states/conformations, depicting lipid accessibility during transient structural changes. Direct cholesterol-GlyR interactions have been identified in different allosteric conformations that can help refine GlyR crystallographic/cryo-electron microscopic models, principally in unresolved regions, and provide insight into ion channel dynamics of gating and desensitization.

## Materials and Methods

### Purification of GlyR from Sf9 insect cells into mixed detergent micelles

Purified GlyR in mixed detergent/lipid micelles were isolated as previously described^67^. Briefly, wild-type (WT) human α1 GlyR was overexpressed in a baculovirus infected *Sf*9 insect cells. Three days post-infection *Sf*9 insect cells were gently pelleted, washed with phosphate buffered saline (137 mM NaCl, 2.7 mM KCl, 10 mM Na2HPO4, 1.8 mM KH2PO4, pH 7.4), and resuspended in a hypotonic solution (5 mM Tris pH 8.0, 5 mM ethylenediaminetetraacetic acid (EDTA), 5 mM ethylene glycol tetraacetic acid (EGTA), 10 mM dithiothreitol (DTT)). An anti-proteolytic cocktail (1.6 µunits/mL aprotinin, 100 µM phenylmethanesulfonyl fluoride, 1 mM benzamidine, 100 µM benzethonium chloride) was added to reduce protein degradation immediately preceding lysis. Cells were lysed by sonication followed by centrifugation (387,000 x g for 30 min) to isolate cell membranes containing GlyR. Cell membranes were washed with a resuspension buffer (hypotonic solution above with 300 µM NaCl) followed by centrifugation again to remove peripheral membrane proteins. The protein pellet was solubilized in 10:1 digitonin:deoxycholate buffer (12 mM mixed lipids (9:1 plant extract (∼95% phosphatidycholine purity, Avanti): egg extract (∼60% phosphatidycholine purity, Avanti) at 1.5 mg/mL, stored as suspended vesicles), 0.10 % deoxycholate, 1.0 % digitonin, 25 mM potassium phosphate (KP_*i*_, pH 7.4), 1 M KCl, 5 mM EDTA, 5 mM EGTA, 10 mM DTT, anti-proteolytic cocktail) overnight, and solubilized micelles were isolated after centrifugation (387,000 x g for 1 hr). GlyR/lipid/detergent micelles were affinity purified on 2-aminostrychnine agarose and eluted competitively with the addition of excess glycine (2M for WT preps) or strychnine-sulfate pentahydrate (1.5 mM for F207A/A288G preps) to the solubilization buffer.

### Reconstitution of GlyR into lipid vesicles incorporating azi-cholesterol

GlyR vesicle reconstitution was completed as previously described^68^ with the following modifications to the lipid composition. All steps were conducted in the dark and at 5°C unless noted. Mixed lipids (9:1 plant extract (95% phosphatidycholine purity): egg extract (60% phosphatidycholine purity) at 15 mg/mL, stored as suspended vesicles) were added to purified GlyR/lipid/detergent micelles to yield a final concentration of 1.5 mg/mL. Cholesterol (15.07 mM in methanol) with 6 µM azi-cholesterol was included to yield >40 mol percent. As previously described, samples were added to dialysis cassette (3500 MW cutoff, Thermo) for dialysis against excess 6.25 mM KP_*i*_ buffer at pH 7.4. The final reconstituted protein pellet was dissolved in 25 mM KP_*i*_ buffer at pH 7.4. GlyR concentrations were quantitated using a modified Lowry assay^69^.

### Photo-crosslinking of azi-cholesterol to GlyR and separation of crosslinked GlyR

Reconstituted GlyR vesicles were placed into quartz cuvettes on ice to maintain temperature. Photocrosslinking of azi-cholesterol to GlyR was completed as previously described^49^ with the following modifications. To enrich for GlyR in the open state, 30 nM ivermectin was added to reconstituted GlyR vesicles (F207A/A288G) immediately before UV light exposure. To enrich for GlyR in the desensitized state, 10 mM glycine was added to reconstituted GlyR (WT) vesicles immediately before UV light exposure. Cuvettes were exposed to a 420 W Hg Arc lamp (Newport, Model 97435-1000-1, 260-320 nm) for 4 sessions of 5 minutes at 7 cm, with 5 minute periods of no exposure in between each UV exposure session to prevent sample warming. SDS-PAGE (11 % resolving, 5 % stacking) separated the crosslinked oligomeric and monomeric GlyR from lipids, with gel plugs excised between migration distances of 250 kDa and 37 kDa, encompassing the mass of oligomeric and monomeric forms of GlyR.

### In-gel Trypsin digestion of crosslinked GlyR

Gel plugs were washed with 50:50 methanol: 50 mM ammonium bicarbonate twice for 40 min with gentle agitation (VWR Thermal Shake Touch, 900 rpm). Gel plugs were dehydrated by adding 500 μL acetonitrile. Once gel plugs turned whitish, acetonitrile was removed and gel plugs were dried in an Eppendorf 5301 Vacufuge Concentrator for approximately 15 minutes. Trypsin solution (10 μL at 20 μg/mL in 50 mM ammonium bicarbonate) was added to gel plugs and incubated on ice for 15 minutes, then incubated overnight at 37°C with gentle agitation (VWR Thermal Shake Touch, 900 rpm). Digested peptides extracted into supernatant and transferred to VWR non-stick microcentrifuge tubes. Tryptic fragments were further extracted by incubating gel plugs twice for 30 minutes in 300 μL of 0.1 % formic acid in 50:50 acetonitrile:H_2_O. The supernatant was collected and combined with initial supernatant. Tryptic extract solution was dried in an Eppendorf 5301 Vacufuge Concentrator.

### Mass fingerprinting of crosslinked cholesterol to GlyR

50:50 Acetonitrile: H_2_O with 0.1 % Formic Acid (50 μL) was added to tubes containing dried tryptic extracts. ESI-Q-TOF-MS measurements were taken using an Agilent 6530 Q-TOF-MS with an Agilent HPLC-Chip II G4240-62006 ProtID-Chip-150, comprised of a 40 nL enrichment column and a 75 µm x 150 mm separation column packed with Zorbax 300SB-C18 5 µm material. The mass spectrometer was run on positive ion mode using internal standards (1221.9906 and 299.2944) for calibration, supplied by Agilent. Mobile phase compositions used were solvent A (95 % H_2_O, 5 % ACN, 0.1 % formic acid) and solvent B (95 % ACN, 5 % H_2_O, 0.1 % formic acid) The nanoflow elution gradient was developed as follows at 0.50 µl/min of solvent A (minute: percent A): 0.00: 95 %, 4.00: 10 %, 6.00: 70 %, 9.00: 50 %, 11.50: 95 %, 13.00: 95 %. Data were processed using Agilent Qualitative Analysis Software 6.0. Cholesterol crosslinked peptides within a 10 ppm accuracy window were identified, accounting for possible peptide modifications (oxidation, acrylamidation).

For MS/MS studies, crosslinked samples were run again under identical conditions on the Agilent 6530 Q-TOF-MS, targeting the specific m/z ratio, charge, and retention time (RT) of the crosslinked peptides identified in MS analysis. Collision-induced dissociation (CID) was used for MS-MS fragmentation following a linear increase in collision energy by m/z using the equation: y=3.7x+2.5 (y= m/z, x= collision energy). CID was performed at ± 0.2 min from initial MS scan RT of each crosslinked precursor ion identified. Data were processed using Agilent Qualitative Analysis Software 6.0 in conjunction with ProteinProspector v5.14.3 available through the University of California, San Francisco.

## Results

Our previous study^49^ identified apo-state azi-cholesterol-α1 GlyR crosslinking as a function of cholesterol concentration (see Figure 1 for structure of azi-cholesterol), where differential crosslinking patterns were observed in comparative studies of negligible cholesterol chemical activity and chemically active cholesterol, suggesting two distinct structural conformations of apo-state GlyR as a function of cholesterol concentration. The dependence of pLGIC activity on cholesterol has been long established and these studies were consistent with FTIR studies demonstrating the uncoupling of ligand binding from pore opening as a consequence of decreased cholesterol content^70^. These studies also provided evidence that CX-MS studies using photoactivatable cholesterol can sensitively probe GlyR conformation. Given the requirement for GlyR activity on more elevated cholesterol concentrations, all of our comparative studies of homopentameric human α1 GlyR examining cholesterol crosslinking as a function of receptor allostery were conducted at the higher cholesterol concentrations at a level consistent with physiological conditions.

**Figure 1.**
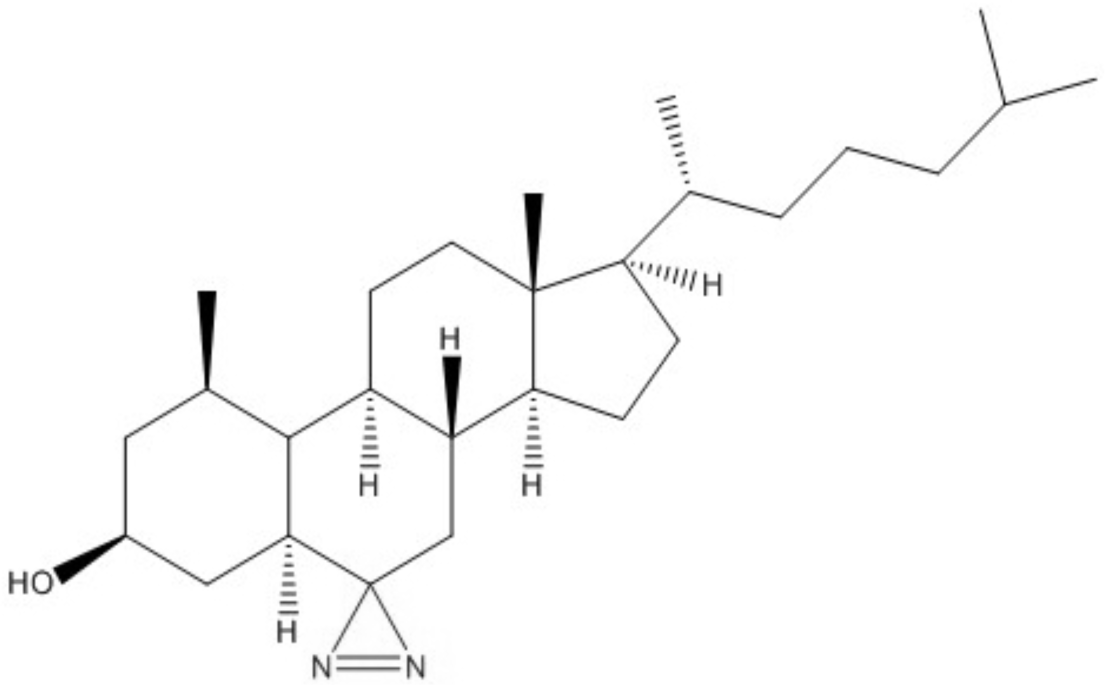
Chemical structure of photoactivatable azi-cholesterol. Chemical structure of cholesterol crosslinker analog, with the reactive site highlighted in blue.

In the previous studies^49^, azi-cholesterol crosslinking of GlyR in its resting state was observed in the pre-M1 extracellular domain on the outer lipid-exposed surface, the extracellular M2-M3 loop, regions of the large intracellular M3-M4 loop, the M4 transmembrane helix, and the post-M4 c-terminal tail (Figure 4A, 5A, Table 1). In this study we extend these studies to examine cholesterol photocrosslinking in a state-dependent manner, conducting comparative crosslinking studies on purified and reconstituted human α1 GlyR enriched in either its resting, open or desensitized states. Ivermectin promotes predominant stabilization of channels in a conducting conformation (open state) in F207A/A288G a1 GlyR^65^. In our hands, F207A/A288G GlyR expressed in insect cells were gated by ivermectin in whole cell patch clamp studies and showed no evidence of desensitization (Tomcho *et al*., manuscript in preparation), consistent with published observations^65^. Under excess ivermectin conditions (1.5 mM) these non-desensitizing channels are expected transition between open or resting states, and comparative CX-MS studies with apo studies should identify mass-shifted ions unique to each study, thus allowing identification of unique cholesterol binding sites restricted to each of these states (or exposed during structural transitions between the resting and open state). Similarly, in order to examine cholesterol accessibility in the desensitized state of the receptor, cholesterol interactions were examined in conditions of excess glycine enriching GlyR in a higher-affinity ligand-bound desensitized conformation. Given that the receptor is expected to exist in resting, open, and (primarily) desensitized states, comparative studies are conducted to identify crosslinking sites uniquely observed in the presence of excess glycine and not in our resting or open state studies.

**Table 1.**
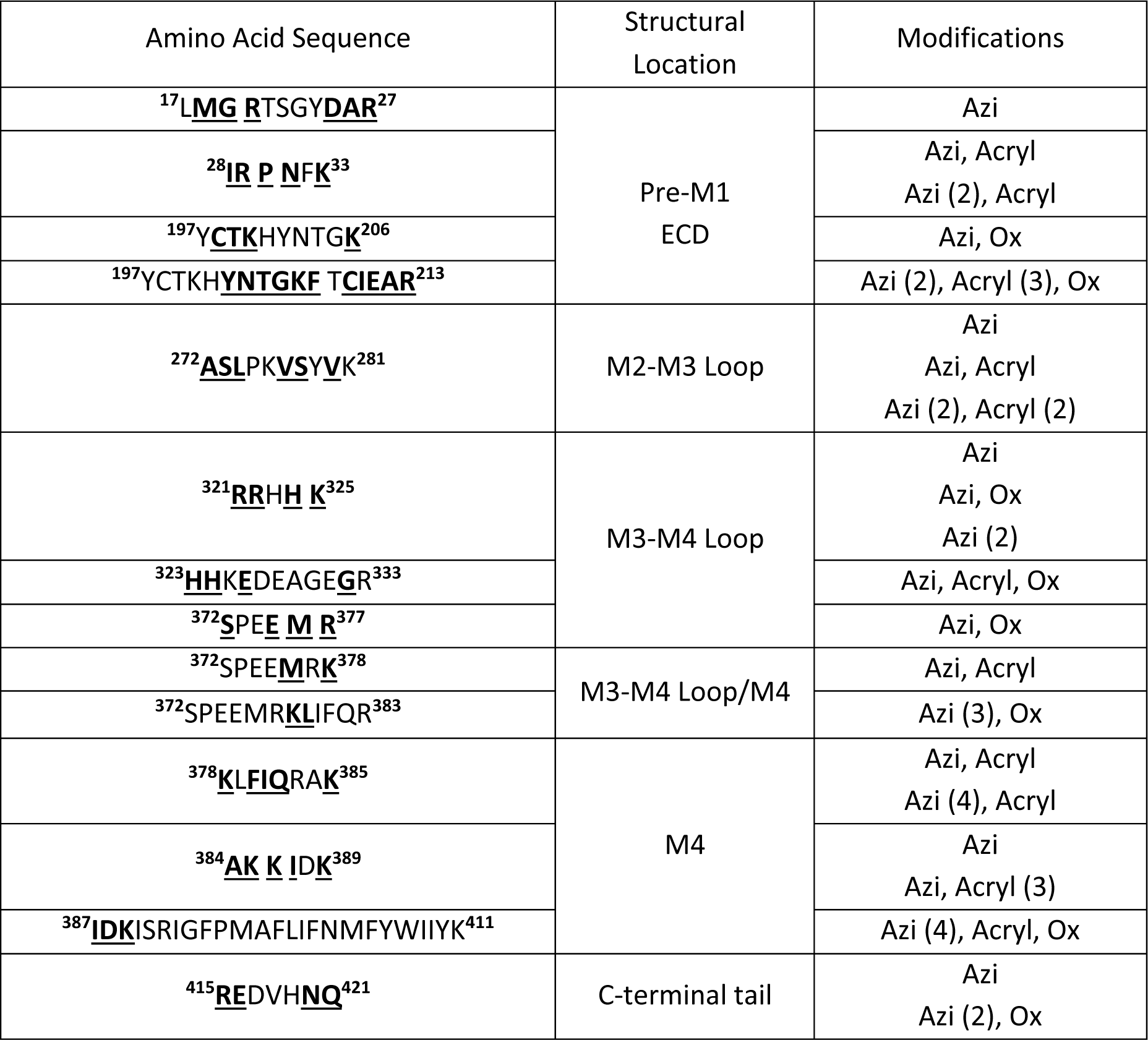
Identified resting precursor/product ion crosslinked peptides at >40 percent cholesterol (from previous study^49^). Identified mass-shifted precursor ions crosslinked with cholesterol (within 10 ppm error, identified in at least 2 of 3 trials) at >40 mol percent cholesterol conditions shown in left column. Sites of covalent modification identified upon analyses of product ions upon CID fragmentation are bolded and underlined; spaces separating amino acids represent single point amino acid crosslinking sites in succession. *Modifications of the precursor ion, including crosslinked cholesterol. Precursor ions identified with different combinations of modifications are shown for each precursor ion. Parenthesized numbers following “Azi” represent the number of crosslinked cholesterol(s) within the mass-shifted peptide.

**Figure 2.**
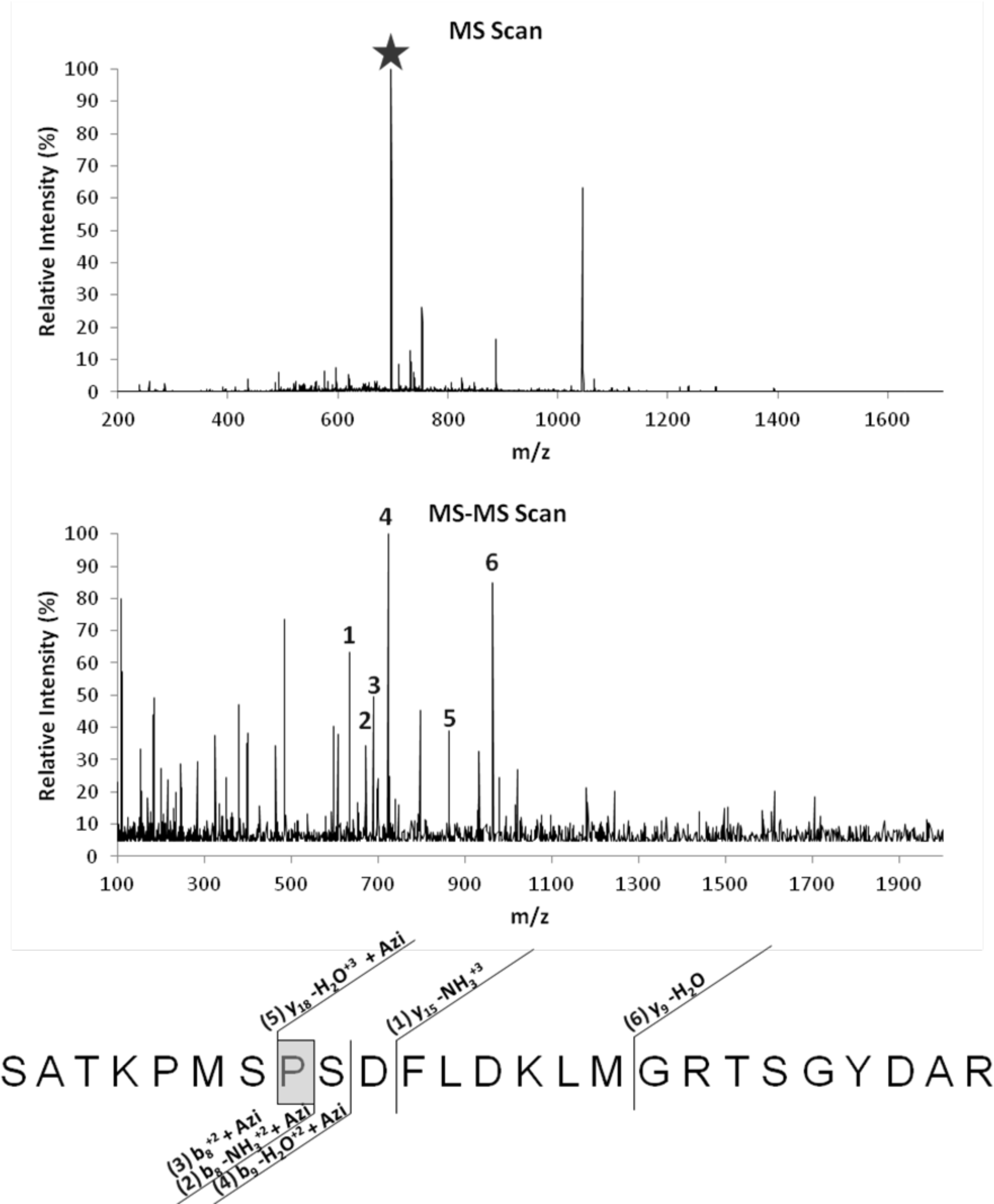
Representative MS-MS analysis. (Top) Representative MS scan identifying crosslinked precursor ion, highlighted with red star. (Middle and Bottom) CID-induced fragmentation of crosslinked precursor ion. The identified product ion fragments (1-6, Middle), including those mass shifted (+ Azi) are highlighted and mapped to peptide to refine crosslinking location (green, Bottom).

**Figure 3.**
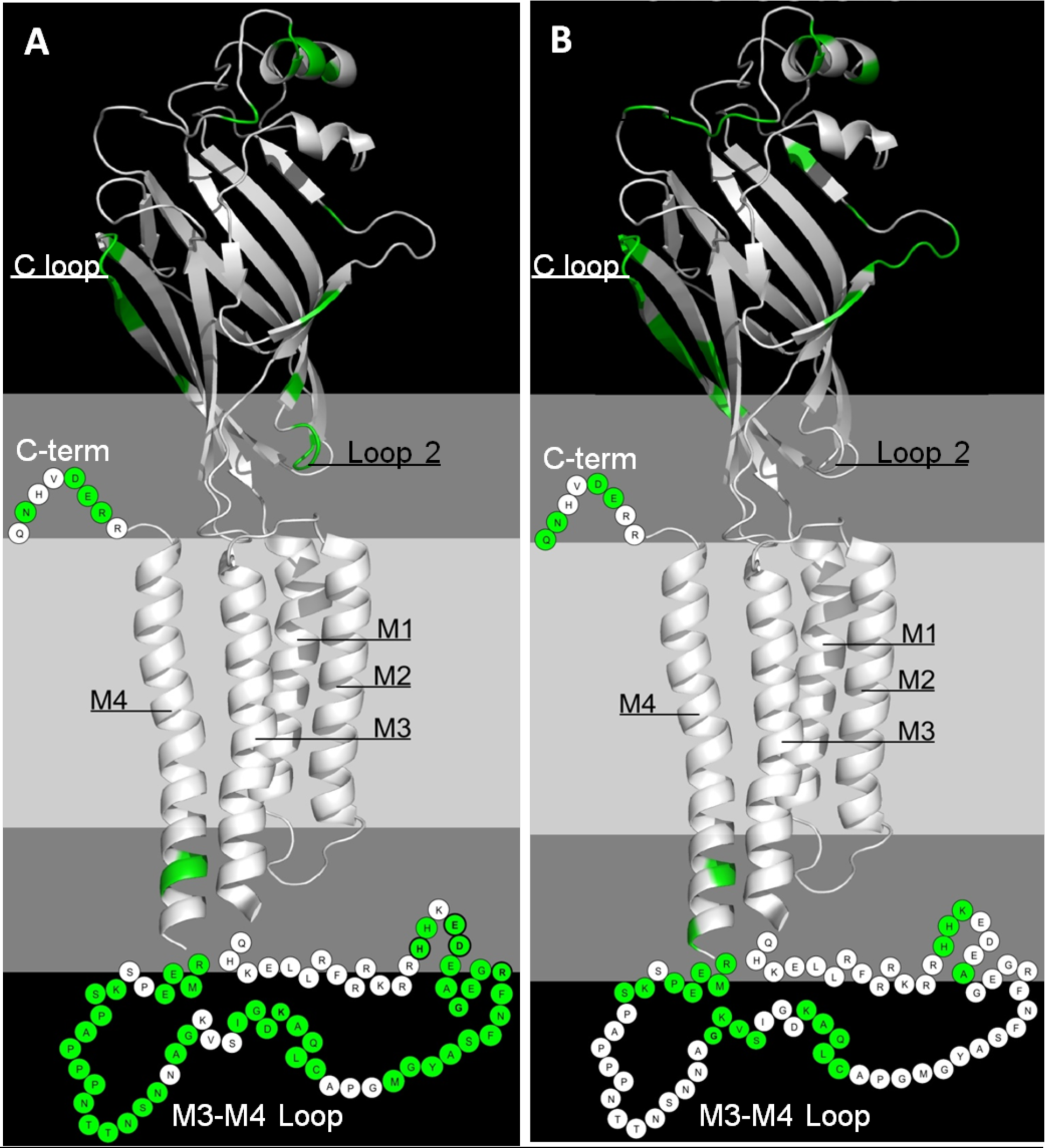
State-dependent GlyR-cholesterol interactions at active cholesterol conditions. Azi-cholesterol crosslinking at (A) open state and (B) desensitized state mapped to a single α1 subunit of zebrafish GlyR (PDB #3JAD) using PyMOL v1.8.^71^ Areas corresponding to the bilayer are shown in gray (∼15Å interfacial regions) and light gray (∼30 Å hydrophobic acyl chain region). Sites of crosslinking in the open and desensitized (bolded and underlined amino acids in Table 2-3, respectively) identified are shown in green. Beads represent regions not resolved in the zebrafish structure (M3-M4 loop and C-terminal tail) with crosslinks arbitrarily placed in close proximity to the lipid bilayer. Bolded beads represent refined sites of attachment.

**Figure 4.**
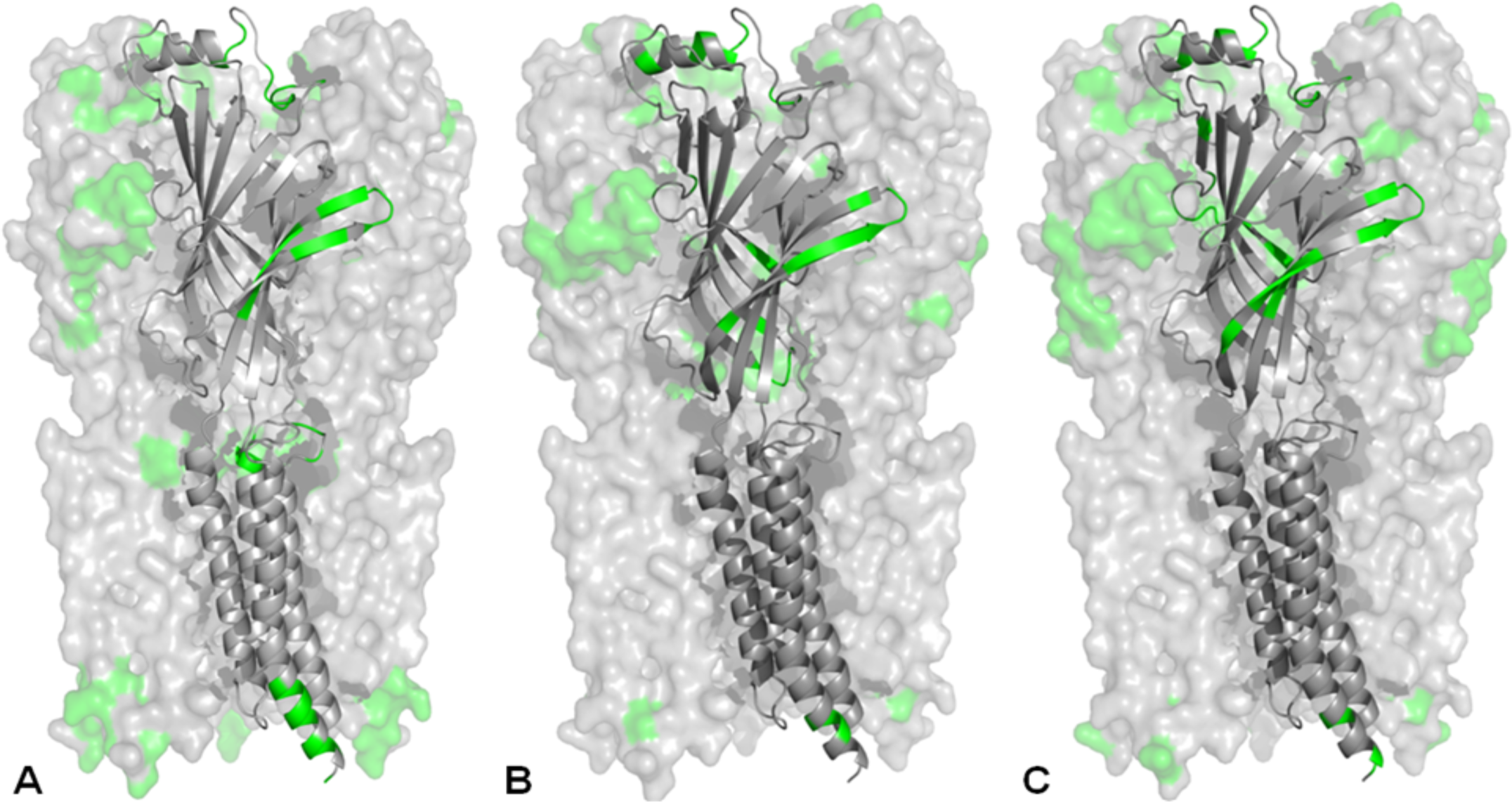
Allosteric GlyR-cholesterol interactions at active cholesterol conditions. Side view of the space-filling GlyR pentamer (with one single subunit shown as ribbon diagram, PDB #3JAD^71^) with sites of crosslinking highlighted in green for the (A) resting state^49^, (B) open state, and (C) desensitized at >40 mol percent cholesterol. Unresolved regions of the receptor are not shown in panels A-C.

Crosslinks were identified by LC-MS/MS in each of the three comparative studies, and a more detailed interpretation of these data is deferred to the Discussion. In all cases, crosslinks were identified in at least 2 of 3 independent sample preparations/MS analysis pairings with peptide identified sequence coverage of GlyR up to 60 percent. Crosslinking analysis was not limited to a single cholesterol crosslinking event, but allowed up to 4 cholesterol crosslinks per tryptic peptide, as cholesterol is distributed in both leaflets of the lipid bilayer having the potential to interact at multiple sites within a single tryptic peptide, as well as “piggybacking” (cholesterol crosslinking to a GlyR bound cholesterol). Identified crosslinks from independent preparations were obtained from a single LC-MS and LC-MS/MS paired experiment. For LC-MS/MS analysis, mass errors in precursor/product ion identification were restricted to < 10 ppm/0.1 Dalton, respectively. Due to the presence of isobaric species (i.e., cholesterol binding at more than one location, such as adjacent amino acids, within a given proteolytic fragment and/or potential piggybacking of cholesterols), strict pairing of the retention times of product ions and their respective precursors allows multiple crosslinks to be identified from isobaric precursor ions in single trials due to their respective differential retention times on the LC-MS/MS platform. A representative precursor/product ion pair is shown in Figure 2.

In studies conducted on non-desensitizing F207A/A288G GlyR, cholesterol crosslinking was identified in the pre-M1 extracellular domain in regions distant of the membrane and closer proximity with the ECD-TMD interface, the entire span of the large intracellular M3-M4 loop, the lower portion of the M4 transmembrane helix, and the post-M4 C-terminal tail (Table 3), and these sites were visualized on a single α1 subunit of zebrafish GlyR (PDB #3JAD)(Figure 3A)^71^. Many crosslinks were found in unresolved regions of the α1 GlyR model (the M3-M4 loop and C-terminal tail), and are depicted as colored beads. These crosslinking events suggest that these sites are in close proximity hydrophobic core of the lipid bilayer, suggesting an intimate association of large swaths of the heretofore unresolved M3-M4 linker with the periphery of the hydrophobic core of the bilayer (the expected depth of the diazirene moiety on azi-cholesterol (Fig. 1). A profile side view of crosslinked regions (Figure 4B) displays these annular associations, yet also exposes more buried non-annular cholesterol interactions in hydrophobic pockets between subunits consistent with nAChR^72^ and GABA_A_R^13^ studies. In bottom-up views of the receptor (Figure 5B) crosslinking locations are localized on outer surface of GlyR, highlighting the expected predominant annular lipid surface accessibility. Cholesterol crosslinking only identified in open state studies, not in the apo-state is suggested to be unique cholesterol-GlyR interactions of the open state. Cholesterol crosslinking unique to the open state GlyR was observed in the pre-M_1_ extracellular domain in regions distant to (residue numbers 3, 10, 15-16, 52-55, 58) and nearing (residue numbers 105-106, 116, 193, 197, 201) the membrane, the large intracellular M3-M4 loop (residue numbers 227-231, 233, 237-241, 245-247, 249, 334-337, 345-348, 350-352, 356-357, 359-371, 374), the M4 transmembrane helix (residue number 383), and the post-M4 c-terminal tail (residue number 417).

**Table 2.**
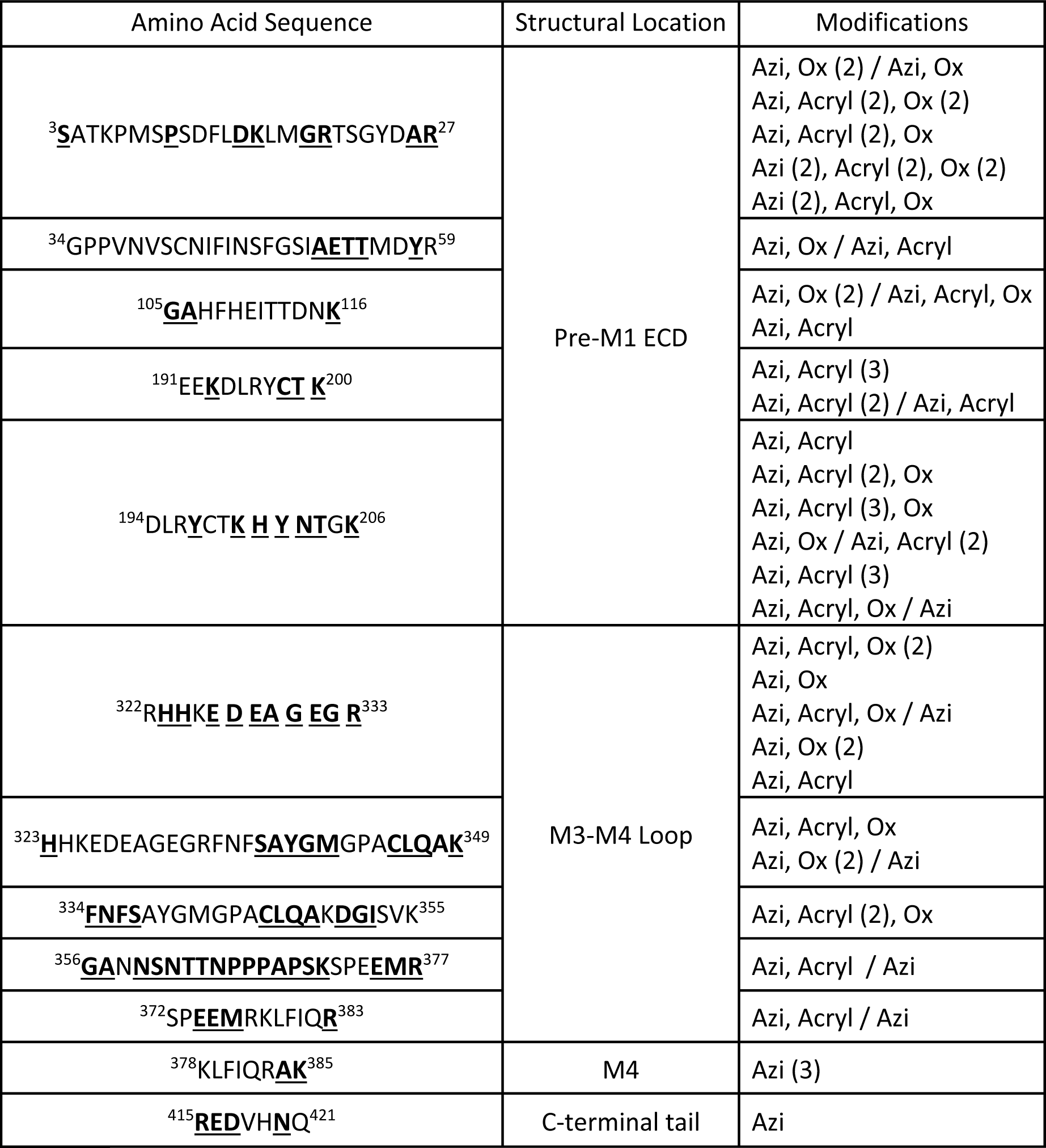
Identified open state precursor/product ion crosslinked peptides at >40 percent cholesterol with conditions as described in Table 1.

**Table 3.**
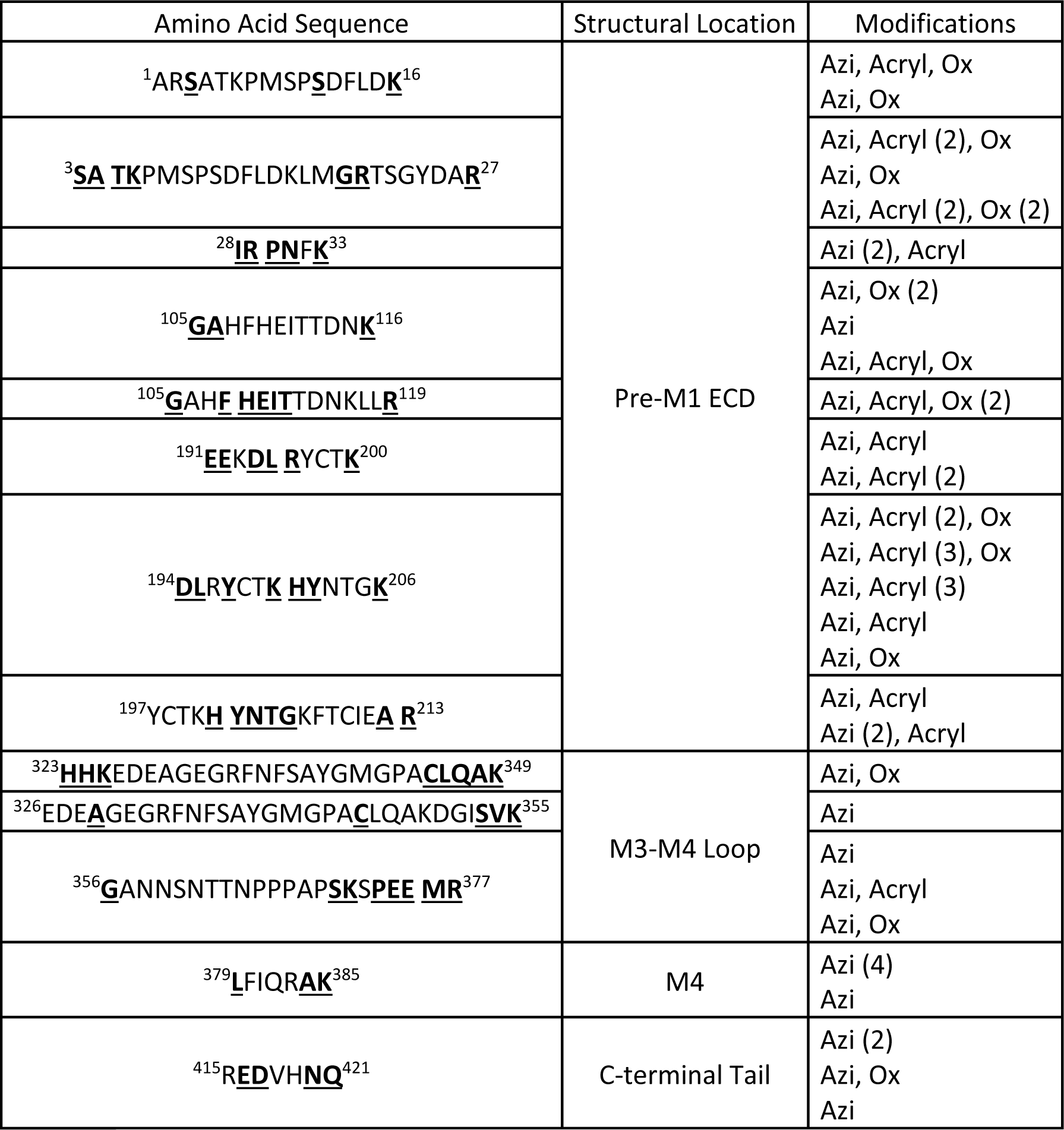
Identified desensitized state precursor ion crosslinked peptides at >40 percent cholesterol with conditions as described in Table 1.

**Figure 5.**
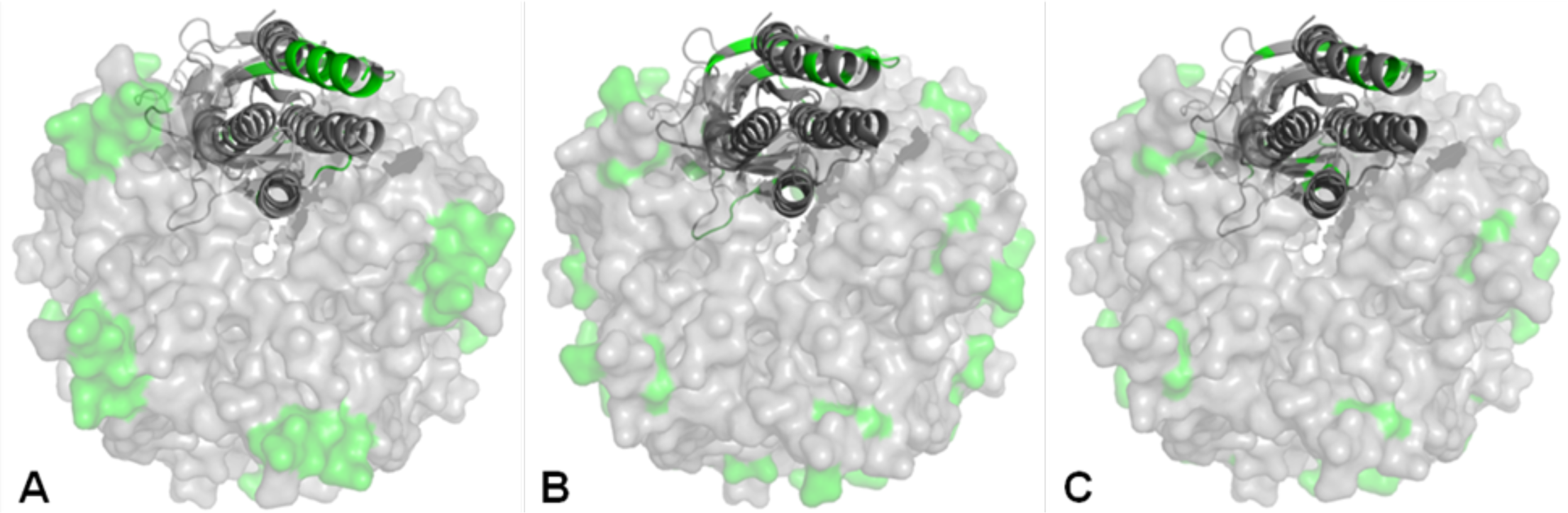
Allosteric GlyR-cholesterol interactions at active cholesterol conditions. Bottom-up view of the space-filling GlyR pentamer (with one single subunit shown as ribbon diagram, PDB #3JAD^71^) with sites of crosslinking highlighted in green for the (A) resting state^49^, (B) open state, and (C) desensitized at >40 mol percent cholesterol. Unresolved regions of the receptor are not shown in panels A-C.

In studies conducted on WT receptor in the presence of excess glycine, cholesterol crosslinking was identified in the pre-M1 extracellular domain in regions distant of the membrane and closer proximity with the ECD-TMD interface, three distinct regions of the large intracellular M3-M4 loop, the lower region of the M4 transmembrane helix, and the post-M4 C-terminal tail, mapped to a single α1 subunit of zebrafish GlyR (PDB #3JAD) (Table 3, Figure 3B)^71^. Crosslinking was observed in unresolved M3-M4 loop and C-terminal tail of the α1 GlyR model (Figure 3B, M3-M4 loop and C-terminal tail, depicted as colored beads) and suggested to be intimately associated with the lipid membrane. Examination of the accessible surface of the pentamer, either by profile side view (Figure 4C) or viewed intracellularly facing outward (Figure 5C), show sites of crosslinking on both the outer surface, annular lipid-accessible regions of GlyR as well as more buried non-annular regions consistent with crosslinking observed in the resting and open state trials. Cholesterol crosslinking uniquely observed in studies in excess glycine are proposed to be unique cholesterol-GlyR interactions of the desensitized state, and was identified in the pre-M1 extracellular domain in regions distant to (residue numbers 4-6, 11) and nearing (residue numbers 108-112, 119, 191-192, 194-196) the membrane, and intracellular M3-M4 loop (residue numbers 353-355, 373).

## Discussion

The differential cholesterol crosslinking patterns discerned in this study between conditions stabilizing the resting, open, and desensitized states of GlyR shows that CX-MS can identify unique cholesterol interactions in a state-dependent manner. This change in the pattern of cholesterol crosslinking between functional states details structural movements between allosteric conformations and define lipid-accessible hydrophobic regions of GlyR, as well as hydrophobic pockets in the receptor. Currently there are structures of GlyR bound to agonist/antagonist^71^, ivermectin^71,73^, and analgesic potentiators^74^, as well as forthcoming cryo-EM structures^63^. However, many of these structures have deletions and mutations for thermostability, are bound to ligand, and often lack the presence of a bilayer, providing limitations in discriminating the entirety of the receptor and understanding allostery of pLGICs. These limitations emphasize the need for continued and improved methodologies to provide additional information for each allosteric state for full-length receptors in the presence of a membrane. State-dependent comparative CX-MS studies offer the unique capability of complementing other high-resolution methodologies to help refine dynamic changes of membrane proteins under physiological conditions. Under such conditions, CX-MS can differentiate subtle structural movements throughout the majority of the protein, including regions unresolved in current structures. The synergistic utility of CX-MS with common structural techniques (x-ray crystallography and cryoEM) can drastically enhance the allosteric understanding of membrane proteins.

In the absence of quantification, this comparative state-dependent cholesterol-GlyR study is unable to distinguish high frequency from low frequency crosslinks identified. Therefore, it is assumed that crosslinking studies conducted in the presence of ivermectin (labeled as open state crosslinking) will capture crosslinks of both resting and open state channels as well as accessibility during state transitions as we are unable to distinguish crosslinking specific to the resting, open, or intermediate structures. Similarly, crosslinking studies conducted in the presence of glycine (labeled as desensitized state crosslinking) will capture crosslinks of the resting, open, and desensitized channels as well as accessibility during state transitions as we are unable to distinguish crosslinking specific to the resting, open, desensitized, or intermediate structures. Unique crosslinks identified within each three conditions tested are suggested to be distinctive cholesterol-GlyR interaction profiles and consist of up to ∼58% of crosslinks identified within each allosteric state. The relative abundance of identified mass ions reflects their ionization, and not their concentration, Given the lack of quantification, no discrimination is made between specific and non-specific sites of cholesterol interaction. All sites of interaction identified via covalent crosslinking are interpreted as stochastic sites of accessibility.

Sites of cholesterol crosslinking determined in CX-MS studies were visualized through mapping on the α1 GlyR strychnine-bound model^71^. This model was selected to most closely resemble the apo-state conditions tested in the previous study given the three available α1 GlyR models (strychnine, glycine, and glycine/ivermectin bound)^71^. To maintain consistency, all crosslinks of this CX-MS study are mapped on this single static model to enhance identification of regions exhibiting changes in lipid accessibility, thereby identifying movements of the receptor underlying gating and desensitization. Consequently, the mapping shown for each allosteric state (open and desensitized) may not accurately represent structural-based crosslink localization identified for each state, however gives an approximate depiction of the crosslinked regions. Exact structural changes based upon state-dependent cholesterol crosslinking patterns mapped on the model are suggested to be a result of either cholesterol or protein relocation. Changes in cholesterol’s interaction location based upon lipid relocation are suggested to be from repositioning within hydrophobic pockets or regions of lipid accessibility. Changes in cholesterol’s interaction location based upon GlyR repositioning causes mapped crosslinked site(s) to differ from state to state, where the directional movement of cholesterol location is the opposite of GlyR actual repositioning.

Shifts of crosslinking patterns within M4 (Figure 6A-C) are proposed to be a result of M4 helix repositioning during gating and desensitization. Comparative CX-MS studies suggest the M4 helix undergoes a twisting during channel activation followed by an outward bending during desensitization. This proposed mechanism reflects the loss of crosslinking from the outer-most lipid accessible side of the M4 helix (resting state residues 378-382, 384-389, Table 1, Figure 6A) to more pronounced and concentrated crosslinking on the inner-most pore-facing side of M4 (open state residues 383-385, Figure 6B) indicating a clockwise twist (top-down view) of the M4 helix. Transition to desensitized state (residues 379, 384-385, Figure 6C) displays an increase in crosslinking lower on M4 at residues closer towards the intracellular domain signifying the outward tilt or translation of the bottom portion of M4 helix. Taken together, state-dependent crosslinking of M4 suggests a outward twisting motion as the helix allosteric transitions which is consistent with general TMD movements observed between cryo-EM GlyR structures^71^. Cholesterol crosslinking within the portion of M4 nearing the M3-M4 loop (bilayer lower leaflet region) is consistent with similar studies of nAChR^56^ showing N-terminal M4 cholesterol crosslinking. Our state-dependent cholesterol CX-MS study expands upon the lipid-channel studies by being able to not only highlight the specific crosslinking locations throughout the entire receptor, yet also distinguish the differential crosslinks in a state-dependent manner.

**Figure 6.**
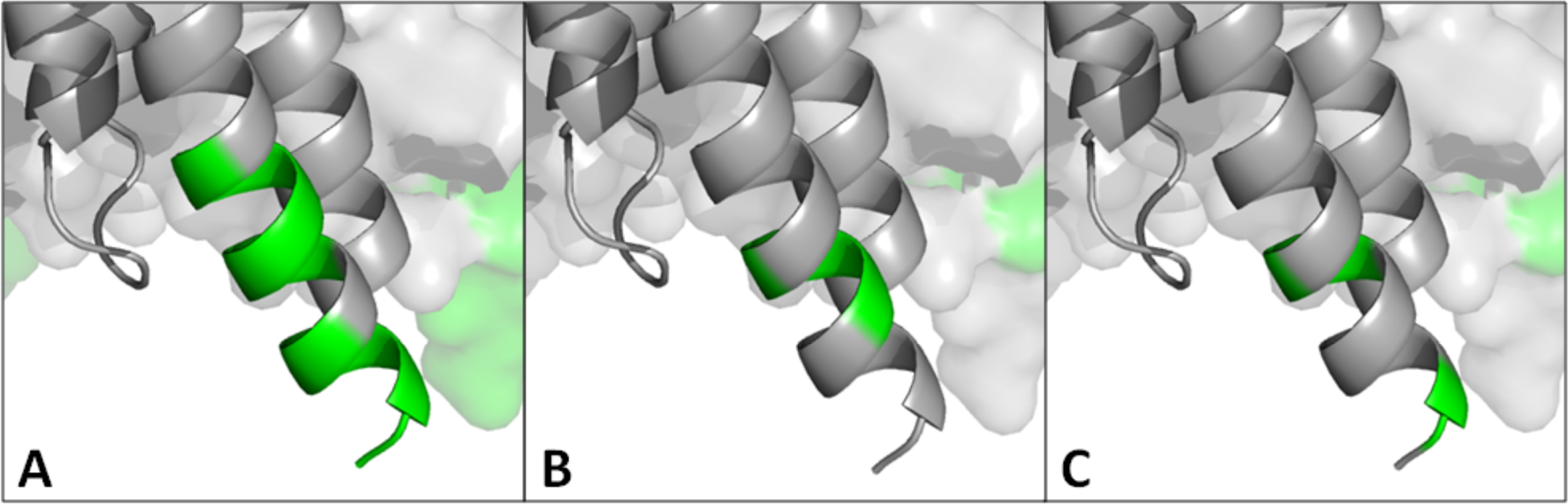
M4 differential cholesterol crosslinking. Comparison of state-dependent cholesterol crosslinking for the resting (A), open (B), and desensitized (C) conditions at > 40 mol percent cholesterol within the M4 helix.

Differential crosslinking patterns were also observed in the ECD nearing the transmembrane domain. In initial CX-MS studies examining cholesterol crosslinking as a function of cholesterol contnet^49^, cholesterol crosslinking in the upper ECD is suggested to arise from azi-cholesterol accessibility to hydrophobic cavities during the detergent-based reconstitution following purification, and these crosslinked regions corresponded to predicted cholesterol interaction locations. In resting state studies (Table 1, Figure 7A), the M2-M3 loop as well as the “C loop” region of the outer β-sheet^71^ flanking above the characteristic Cys-loop were extensively labeled, suggesting the incorporation and presence of cholesterol into the upper leaflet of the bilayer along with a hydrophobic pocket between neighboring subunits. The crosslinked M2-M3 loop residues adjoin a hydrophobic pocket at the ECD-TMD core^75^ in a region shown to harbor transmembrane cavities that bind cholesterol^13^, anesthetics^76^, and correspond to predicted cholesterol recognition motifs identified in proteins^49^. Cholesterol crosslinking within and near the C Loop encompass the hydrophobic β-core conserved among pLGICs^77^ correlating with determined CRAC/CARC interaction motifs^49^. This is a region of dynamic structural alterations during channel gating and desensitization^78^ as well as Zn^2+^ modulation^79^. Comparing apo-state studies with conditions stabilizing the open state (Figure 7B) reveals a loss of M2-M3 loop crosslinking (residues 272-274, 277-278, 280, Table 1) in conjunction with a shift of crosslinking from exclusively outer β-sheet C Loop region to incorporate regions within the inner β-sheet (residues 52-55, 58, 105-106, 116) and lower “Loop 2”^80^ that adjoin ECD-TMD interface, suggesting a reorientation of the ECD during gating that either exposes the lower crosslinked residues identified or alters the hydrophobic cavity around/within the β-sheets. Juxtaposing crosslinking in conditions that stabilized an open channel with desensitization (Figure 7C) exposed nearly identical crosslinking with the exception of the loss of crosslinking in the lower Loop 2 (residues 52-55, 58), suggesting allosteric repositioning of the lower ECD-upper TMD interface during desensitization. Taken together, the ECD structure appears to be more similar between conducting and desensitized GlyRs than apo-state channels and that more dynamic movements are observed during channel activation in the lower ECD region where expansion from the central pore is noted^12,71^.

**Figure 7.**
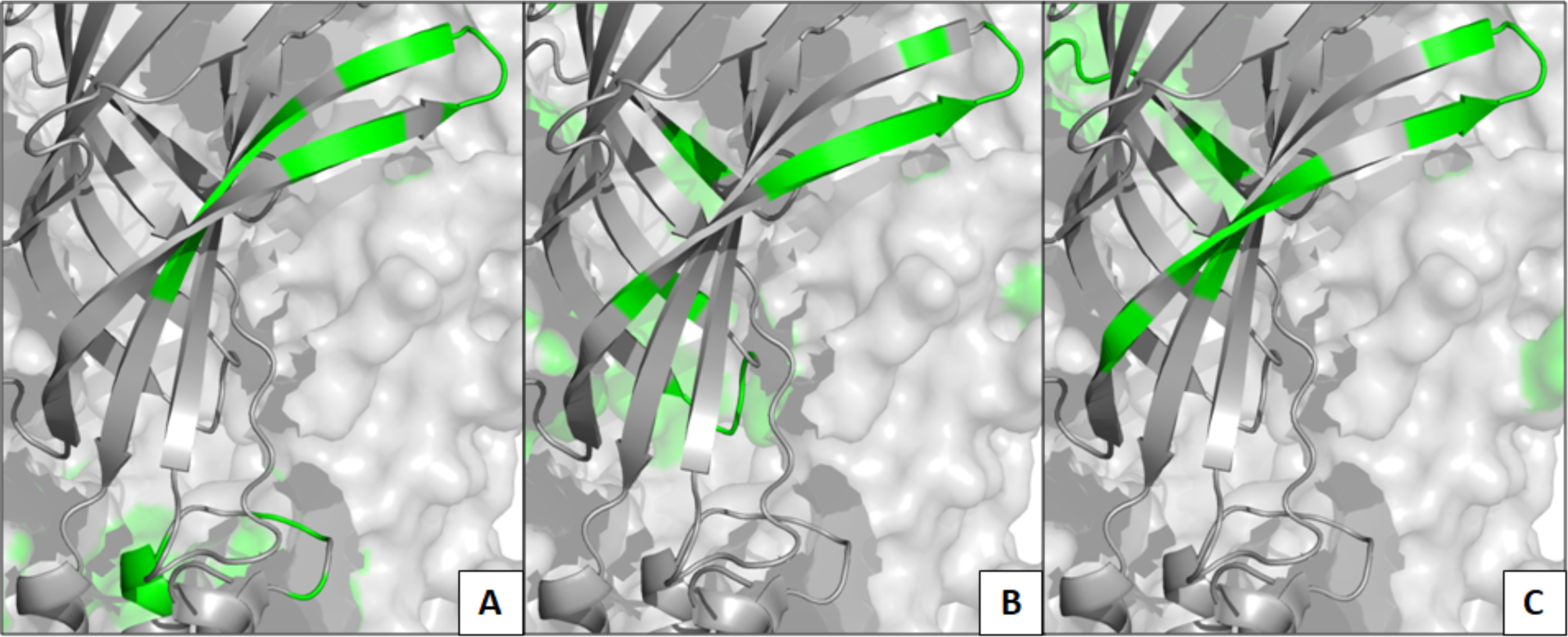
ECD differential cholesterol crosslinking. Comparison of state-dependent cholesterol crosslinking for the resting (A), open (B), and desensitized (C) conditions at > 40 mol percent cholesterol within the pre-M1 ECD region at the transmembrane interface.

The large intracellular M3-M4 loop was highly crosslinked by cholesterol in each allosteric state, providing direct experimental evidence of an intimate association of this loop with the lipid membrane. In addition, the differential patterning of crosslinking as a function of allosteric state indicates that the loop structure changes dramatically as a function of allostery. Contrasting crosslinking patterns from the resting apo-state^49^ to open state. studies (Figure 3A) show nearly identical patterns with the dramatic emergence of crosslinking within central region of the M3-M4 loop (residues 327-341, 345-352, 356-357, 359-371) suggesting a reorientation of the M3-M4 loop during the gating process exposing previously lipid-inaccessible residues. Allosteric shifting from open to desensitized states illustrates the retention of cholesterol crosslinking at the ends (observed ubiquitously) and middle of the M3-M4 loop (Figure 3A-B), suggesting minimal conformational rearrangement that promotes less lipid accessibility. Taken together, the cholesterol crosslinking suggests three distinct allosteric conformations of the M3-M4 loop most notably through the emergence of profound crosslinking in open and desensitized states exposing novel evidence pertaining to this elusive region of GlyR. Given that the M3-M4 loop is often truncated or is unresolved in the reported structures of pLGICs^71,73,81,82^ these results are amongst the first to examine structural transitions of this loop in the context of the functional full-length receptor. Given that this loop is involved in binding to other intracellular proteins and is post-translationally modification^83,84^ these effects may be partially due to altered interactions with lipids/accessibility or influenced by the lipid’s modulation of structure.

In a similar manner, the unresolved post-M4 C-terminal tail of GlyR absent in structural models^71^ was crosslinked by cholesterol in all three allosteric states assayed (Figure. 3 and Figure 1 from previous publication^49^) suggesting that some residues within this region are also intimately associated with the lipid membrane. This area of the receptor is highly dynamic upon activation/desensitization and is implicated in Zn^2+^ modulation^79^. Cholesterol crosslinking between allosteric states was nearly identical in the C-terminal tail, with subtle shifts in cholesterol labeling indicating no major differences (residues 415-417, 420-421). This suggests minimal changes in lipid accessibility of the post-M4 c-terminal tail of GlyR during receptor activity.

Taken together, cholesterol associates and intimately interacts with α1 GlyR in all domains. Differential crosslinking patterns of cholesterol were observed as a function of allosteric state highlighting changes in cholesterol accessibility as a function of GlyR allostery. Differential crosslinking patterns in the ECD, ECD-TMD interface, M3-M4 loop, and M4 highlight conformational changes during gating and desensitization. Thus, CX-MS sensitively provides specific and accurate structural information in a state dependent manner without potential detrimental structural alterations, providing a complementary approach to other high-resolution structure methodologies.

## Acknowledgement

This work was supported by grants from the NIH (R21 MH098127) and a PA Dept. of Health CURE award. The purchase of the mass spectrometer was supported by the NSF (MRIDBI-0821401).

## Notes

### Competing Interest Statement

The authors have declared no competing interest.

